# Deconvolution of Nucleic-acid Length Distributions: A Gel Electrophoresis Analysis Tool and Applications

**DOI:** 10.1101/636936

**Authors:** Riccardo Ziraldo, Massa J. Shoura, Andrew Z. Fire, Stephen D. Levene

## Abstract

Next-generation DNA-sequencing (NGS) technologies, which are designed to streamline the acquisition of massive amounts of sequencing data, are nonetheless dependent on various preparative steps to generate DNA fragments of required concentration, purity, and average size (molecular weight). Current automated electrophoresis systems for DNA- and RNA-sample quality control, such as Agilent’s Bioanalyzer ^®^and TapeStation ^®^ products, are costly to acquire and use; they also provide limited information for samples having broad size distributions. Here we describe a software tool that helps determine the size distribution of DNA fragments in an NGS library, or other DNA sample, based on gel-electrophoretic line profiles. The software, developed as an ImageJ plug-in, allows for straightforward processing of gel images, including lane selection and fitting of univariate functions to intensity distributions. The user selects the option of fitting either discrete profiles in cases where discrete gel bands are visible, or continuous profiles, having multiple bands buried under a single broad peak. The method requires only modest imaging capabilities and is a cost-effective, rigorous alternative characterization method to augment existing techniques for library quality control.

## INTRODUCTION

Next-generation DNA Sequencing (NGS) has rapidly become an indispensible tool in virtually every life-science discipline. NGS workflows entail various DNA-sample manipulations including PCR and enzymatic reactions to prepare DNA fragments of specific concentration, purity, and size in ways that are compatible with a particular sequencing platform (1). The quality of the NGS library has substantial influence on the success of a sequencing run, affecting both sequence validity and the number of reads (2). Many current quality-control (QC) protocols require costly instruments and consumables to assess the library average size, specifically capillary-gel electrophoresis tools such as Agilent’s Bioanalyzer^®^ and TapeStation^®^ products. With the exponential growth of genome-wide analyses and large data sets that hinge on NGS, it is crucial to optimize the accuracy and cost-effectiveness of each workflow step.

Agarose-gel electrophoresis is arguably the most widely used method for separating biopolymer components in a sample based on physical, chemical and topological properties (3–9). As an important example, a commonly used workflow relies on gel electrophoresis to separate DNA fragments generated by Illumina’s Nextera^®^ Tn5 transposition-based tagging/fragmentation protocol. The Nextera^®^ kit produces sequencing libraries through enzymatic shearing of input DNA. Gel electrophoresis uses the molecular-weight dependence of linear DNA’s mobility in polymer gels to separate the resulting fragments along the direction of migration (7, 8). In applications such as typical analyses of restriction digests the products consist of a small number of individual fragments, producing a characteristic pattern of discrete bands. In contrast, a variety of random-shearing-based protocols (including Nextera^®^ and other transposition-based protocols) can be used to produce broad, ideally uniformly distributed, fragment libraries (10). Thus, agarose-gel separations of tagmentation products, because of the large number of possible fragment sizes, generally yield a quasi-continuous distribution of DNA molecules along the path of migration.

QC on the fragment library can greatly improve the sequencing output and reduce both PCR bias and systematic errors (11, 12). Our image-analysis tool is designed to efficiently analyze both discrete and continuous gelelectrophoresis profiles. The technique fits a univariate composite Gaussian function to approximate the profile. Amounts of each species in the sample are then estimated using numerical integration. The goal is to obtain information about the size distribution of fragmented DNA products, from which the average molecular size and bulk concentration of DNA ends can be determined. In the particular case of amplicon-based sequencing such as Illumina’s platforms, accurate knowledge of such fragment-size distributions is essential to optimize cluster density in sequencing protocols.

## MATERIALS AND METHODS

### Library preparation and gel-electrophoresis-based size selection

Human, *C. elegans*, or bacteriophage-lambda genomic DNA (*λ* DNA) were incubated with Nextera XT Tagmentase^®^ (Illumina), an engineered Tn5 transposase, for at least 15 min at 37 °C. DNA fragments underwent PCR-based amplification using Illumina’s Nextera^®^ primers and the resulting libraries were subjected to electrophoresis for the indicated time in 1% agarose gels (Lonza) at 3.0 V cm-1 in TBE buffer (50 mM Tris-borate, 1 mM Na_2_EDTA; pH 8.4). Gels and electrophoresis buffer contained 0.5 µg mL-1 ethidium bromide unless otherwise noted. NGS libraries consisted of tagmented DNA reisolated from excised regions of gel lanes using a Zymoclean Gel DNA Recovery^®^ kit (Zymo Laboratories) (13).

A reference mixture of *λ*-DNA fragments was prepared by digesting *λ* DNA to completion with 2 units of *Aci*I (New England Biolabs) per µg of *λ* DNA (New England Biolabs) for 2 h at 37 °C. A virtual digest performed using SnapGene^®^ showed that the *λ*-*Aci*I digest consists of 517 DNA fragments and 206 discrete fragment sizes distributed between 2 and 1086 bp. In addition to the *λ*-*Aci*I reference mixture, DNA ladders were used as markers: HiLo ladder (Bionexus, Inc) and an NEB 100-base-pair (bp) ladder (New England Biolabs).

### Image acquisition

Several cameras were used to record digital images of the agarose gels. Both high bit-depth (16-bit) grayscale images and RGB (24-bit) color images were acquired. Table 1 lists the different cameras used for the comparison and the specifications of the acquired images.

**Table 1.**
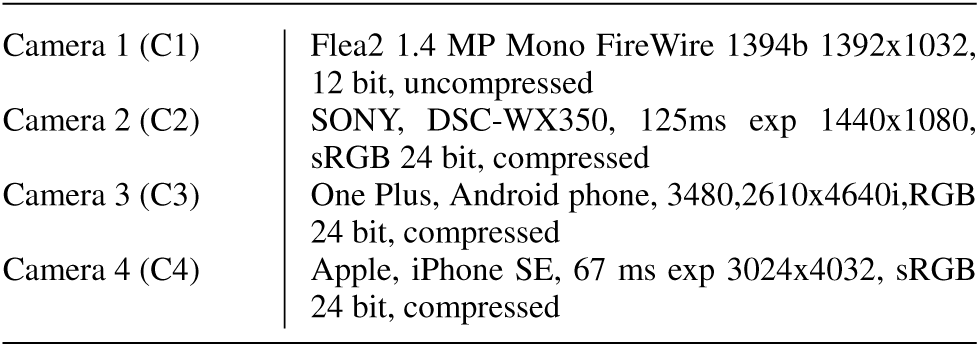
Cameras used for image acquisition

|  |  |
| --- | --- |
| Camera 1 (C1) | Flea2 1.4 MP Mono FireWire 1394b 1392x1032, 12 bit, uncompressed |
| Camera 2 (C2) | SONY, DSC-WX350, 125ms exp 1440x1080, sRGB 24 bit, compressed |
| Camera 3 (C3) | One Plus, Android phone, 3480,2610x4640i,RGB 24 bit, compressed |
| Camera 4 (C4) | Apple, iPhone SE, 67 ms exp 3024x4032, sRGB 24 bit, compressed |

### Sequencing protocol and bioinformatic analysis

Gel-purified DNA libraries were sequenced using the MiSeq^®^ platform (Illumina) according to the vendor’s recommended protocol. Paired-end reads were obtained by sequencing DNA libraries using 150v3 MiSeq^®^ kits (Illumina) for 76 paired cycles. Insert sizes were determined bioinformatically from the distance between paired reads (R1 and R2) after alignment. The size histogram of sequenced reads was obtained as described by Shoura et al. (13).

### Plug-in operation

The workflow for analysis of the gel images is implemented using an ImageJ (https://imagej.net/ImageJ2) plug-in (Gel Lanes Fit). The ImageJ platform was chosen because of its wide distribution as an open-source platform for application and development of image analysis tools. The JFreeChart (http://www.jfree.org/jfreechart) library was used for output plots and the Apache Commons Math (http://commons.apache.org/proper/commons-math/) library was used to implement the fitting algorithm.

The Gel Lanes Fit plug-in includes a number of features that streamline the processing of gel images. Each gel lane is associated with a rectangular *region of interest* (ROI). The dimensions of each ROI can be selected interactively through the plug-ins interface and, as the ROI is resized, the line-profile plot of the corresponding gel lane is dynamically updated. The interface provides access to parameters for non-linear least-squares fitting of the line profile discussed below.

### Function describing the line profile

The image-intensity line profile generated for the corresponding ROI is modeled using a composite univariate function of a coordinate *y* corresponding to the apparent electrophoretic migration distance of DNA fragments in the gel. This function is the sum of a background signal and a set of discrete Gaussian peaks associated with each DNA species in the mixture. In cases where there are relatively few species for which the centers of mass of each component are well separated relative to peak width (defined in terms of full peak width at half-maximum (FWHM), for example), the gel pattern is made up of discrete bands and the line profile consists of discrete peaks. Continuous distributions, in which the separation of adjacent peak maxima is comparable to their respective FWHM values, are expected to have the appearance of a “streak” or “smear.”

Some background contribution to the signal is expected due to scattering from the gel and fluorescence from unbound ethidium dye. The background function maximum is assumed to be less than the smallest minimum corresponding to peaks in the discrete case. A constraint on the maximum degree of the background polynomial and its maximum slope (first derivative) can be set by the user using the plug-ins interface.

Equation **1** shows the target function *F* (*y*). *B*_*q*_ (*y*) is a polynomial of degree *q*. Both *q* and a limit on the first derivative of *B*_*q*_ can be selected by the user. *G*_*i*_ are Gaussian functions with given amplitude, *a*_*i*_, mean, *m*_*i*_, and standard deviation *σ*_*i*_.

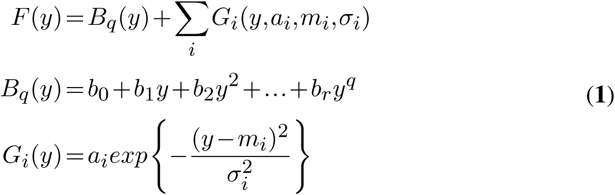

The function is fitted using the Levenberg-Marquardt least-squares algorithm ref. For discrete patterns, each peak is assumed to be Gaussian with the mean at the location of a local maximum and having variance proportional to the band spread. Thus, initial guesses for *m*_*i*_ and *σ*_*i*_ are made using local maxima and peak widths. The peak tolerance parameter sets a threshold for the peak height in the initial guess and thereby filters any peaks contributed by noise. This parameter is defined as a percentage of the full line-profile intensity range.

### Discrete and known continuous patterns

Well-resolved image-intensity line profiles show peaks that correspond to various species present in the sample. However, as discussed above, well-separated peaks cannot always be obtained. The function given in Equation **1** is assumed to be valid for both discrete and continuous profiles, but stronger constraints are needed to fit profiles composed of under-resolved peaks. If a small peak overlaps a larger peak, it cannot be detected automatically. However, an initial guess can be included using the custom peak feature, which specifies the potential location and width of the undetected peak.

When the line profile approaches a continuum, there are no discernible peaks and a different fitting strategy has to be used. Each fragment in a known distribution is used to define constraints on the location and the width of the Gaussian peaks in Equation **1**. As a model for a quasi-continuous distribution of DNA-fragment sizes, we used the mixture of products generated by complete digestion of *λ* DNA by the restriction enzyme *Aci*I. This product mixture contains 517 fragments of known size (see Library preparation and gel-electrophoresis-based size selection) and constitutes a basis set for fitting unknown, quasi-continuous fragment distributions. The vast majority of fragments in this mixture lie in a size range comparable to that selected during library preparation.

An initial guess for the position (mean) of a fragment’s peak is determined from the locations of known species in a discrete, well-resolved ladder of standard fragments of known molecular weight. The position of the unresolved fragments along the continuum is interpolated from the discrete-ladder pattern using the spline function in equation **2**, with spline nodes located along positions that correspond to the maxima of ladder bands. We assume a piecewise linear relationship between the logarithm of the molecular weight, log_10_ (*M*), and displacement in the direction of migration, *y* (9). The molecular weight, *M*, of a particular double-stranded fragment is calculated from the molecule size in base pairs, *l*, and the linear relation *M* (*l*)= 607.4*l* +157.9. Here, 607.4 is the molecular weight of the average nucleotide pair and 157.9 is the combined molecular weight of two 3’- or 5’-monophosphate groups, which are the termini generally left behind by the restriction enzyme (14). The displacement in the direction of migration, *y*, observed in the gel is estimated using Equation **2**, based on a piecewise-linear relationship between log_10_(*M*) and *y* ref. Thus, the linear segments are delimited by the components of the standard ladder of molecular weights, *M*_*ladder*_ = {*M*_1_,*…, M*_*k*_}, and the respective observed displacements, *Y*_*ladder*_ ={*Y*_1_,*…, Y*_*k*_}Each segment of *y* is defined over the intervals *M*_*i-*1_ *< M* ≤*M*_*i*_ for the ladder species *i* = 2, *…, k*.

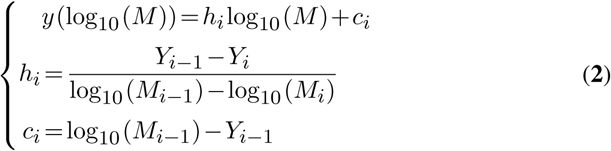

In the case of a *discrete* pattern the standard deviation of the Gaussian peak associated with the band, *σ*_*i*_, is estimated using the FWHM of the peak intensity profile. For *continuous* patterns, a value of *σ*_*i*_ for each fragment in the distribution is determined by interpolation of a piecewise least-squares linear fit to the *σ*_*i*_ values corresponding to bands in a discrete-ladder standard. Additional constraints are applied to the final profile fit by restricting deviations allowed relative to the original *σ*_*i*_ estimates. The observed band intensity, or peak amplitude, *a*_*i*_, is assumed to be proportional to the product of mass fraction and DNA molecular weight. The initial guess for the peak amplitude is thus estimated as *a*_*i*_ = *f*_*i*_*w*_*i*_*d*, where *f*_*i*_ is the fragment’s relative abundance, *w* is the molecular weight normalized by the maximum value, and *d* is the full-scale range of the line profile.

Therefore, for a generic continuous profile with *unknown* fragments, the initial guess is represented by a set of peaks associated with a *known* distribution of fragments and assumed similarities between known and unknown distributions of fragments. The final fit is constrained so that the location and FWHM of each Gaussian component is similar to that of the initial guess with the estimate for the initial amplitude of each component based on the abundance and size of fragments in the initial distribution. Adherence of the amplitude for each peak to that expected from the known distribution is constrained by an area-drift parameter described below.

### Area-drift constraint for continuous line profiles

Initial peak areas based on the parameters corresponding to the known distribution are taken as elements of an array, *P* in Equation **3**, calculated for peak *i* as 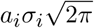, from the peak’s amplitude, *a*_*i*_, and standard deviation, *σ*_*i*_. At each fitting iteration, the set of parameters is updated based on the algorithm’s gradient requirements. The new list of parameters is first checked to exclude non-physical parameter values, for *a*_*i*_, *m*_*i*_ and *σ*_*i*_. After the exclusion of non-physical parameter values, the algorithm recalculates an array of areas, *Q*, in Equation **4**. Particular constraints are enforced so that the peak-area values from the initial guess retain their proportionality, based on the known DNA mass distribution in the *λ*-*Aci*I digest.

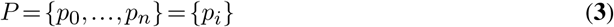

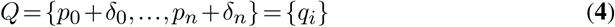

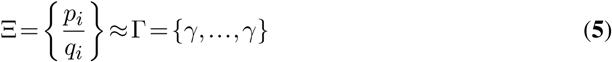

The array Ξ contains element-by-element ratios of *P* and *Q* arrays. The applied area constraint ensures that *Q* stays proportional to *P*. Thus, the elements in the array of ratios, Ξ, are bounded, and would ideally have the same values as those in the array Γ, in Equation **5**. This constraint is enforced using the condition in Equation **6**. If the standard deviation of the elements in Ξ, *s*({*ξ*_*i*_*}*), exceeds a value *E*, chosen by the user, then the array of areas *Q* is adjusted to compensate, using *Q′*.

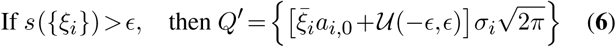

Since the FWHM is subjected to a stricter constraint (see previous subsection), the area constraint is mostly enforced using the peaks’ amplitudes, *N ′* ={*a*_1_,*…, a*_*k*_, which are obtained by scaling the array of initial amplitude guesses, *N* ={*a*_1,0_, *…, a*_*k*,0_}, by 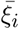, the sum of mean values of the components of {*p*_*i*_*/q*_*i*_*}* plus a uniformly distributed random increment, *𝒰*.

## RESULTS

### Discrete profiles

The major goal of plug-in development was to provide a tool that was equally capable of quantifying discrete, banded, as well as continuous, gel profiles. An example of discrete-band quantification is shown in Figure 1, in which the plug-in is used to analyze a standard ladder composed of known molecular weights. The location of clearly-defined bands is automatically detected using the location of local maxima in the profile. The peak position, maximum intensity above background, integrated intensity, and the FWHM are reported by the plug-in for each peak in the gel lane. As an example of a discrete profile fit, Figure 1 shows the output plot with the fitted profile’s distribution. The plug-in also reports the *root-mean square* (RMS) difference between the fit and the line profile, a measure of the closeness of the fitted function to the original profile (not shown). This discrete-profile fitting strategy applies in general, but it is used specifically by the plug-in to analyze continuous line profiles.

**Figure 1.**
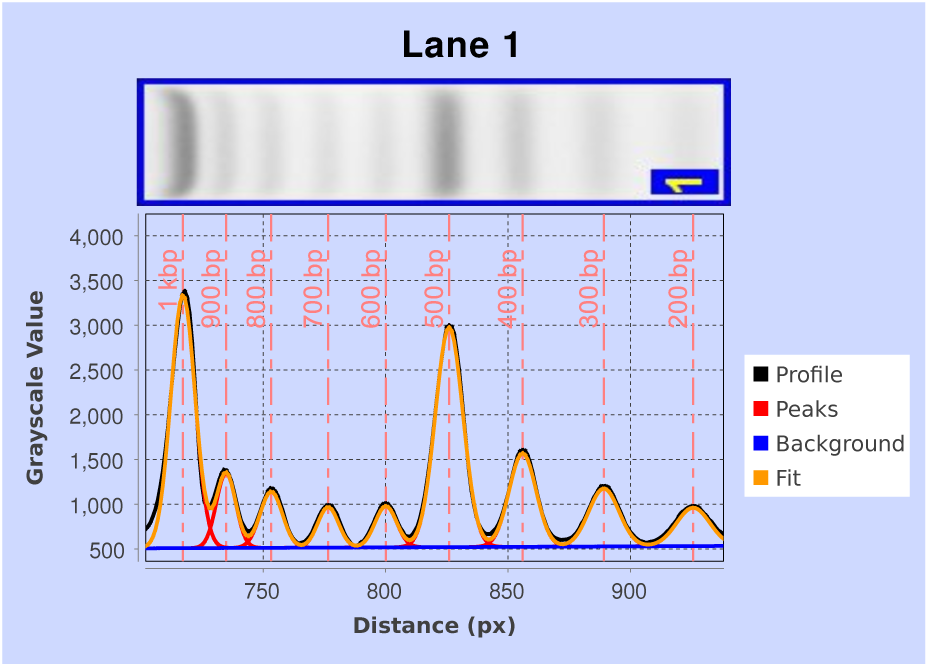
Fit to a discrete profile obtained for gel lane shown in the top panel (also present in lane 1 of Figure 2a). The complete nonlinear least-squares fit (orange) and its component background (blue) and Gaussian peaks (red) are shown. The vertical dashed lines (light red) indicate the locations of peak maxima, corresponding to known molecular-weight values for the ladder species.

### Continuous profiles

The plug-in can quantify profiles with sub-optimally resolved bands, such as DNA-fragmentation products obtained with the Illumina Nextera^®^ tagmentation kit, and quantitate the average size of fragments contained in the mixture. Lanes labeled 2 through 5 in the gel in Figure 2a and 1 through 4 in Figure 2c are all examples of continuous gel profiles, in which single bands are not visible.

**Figure 2.**
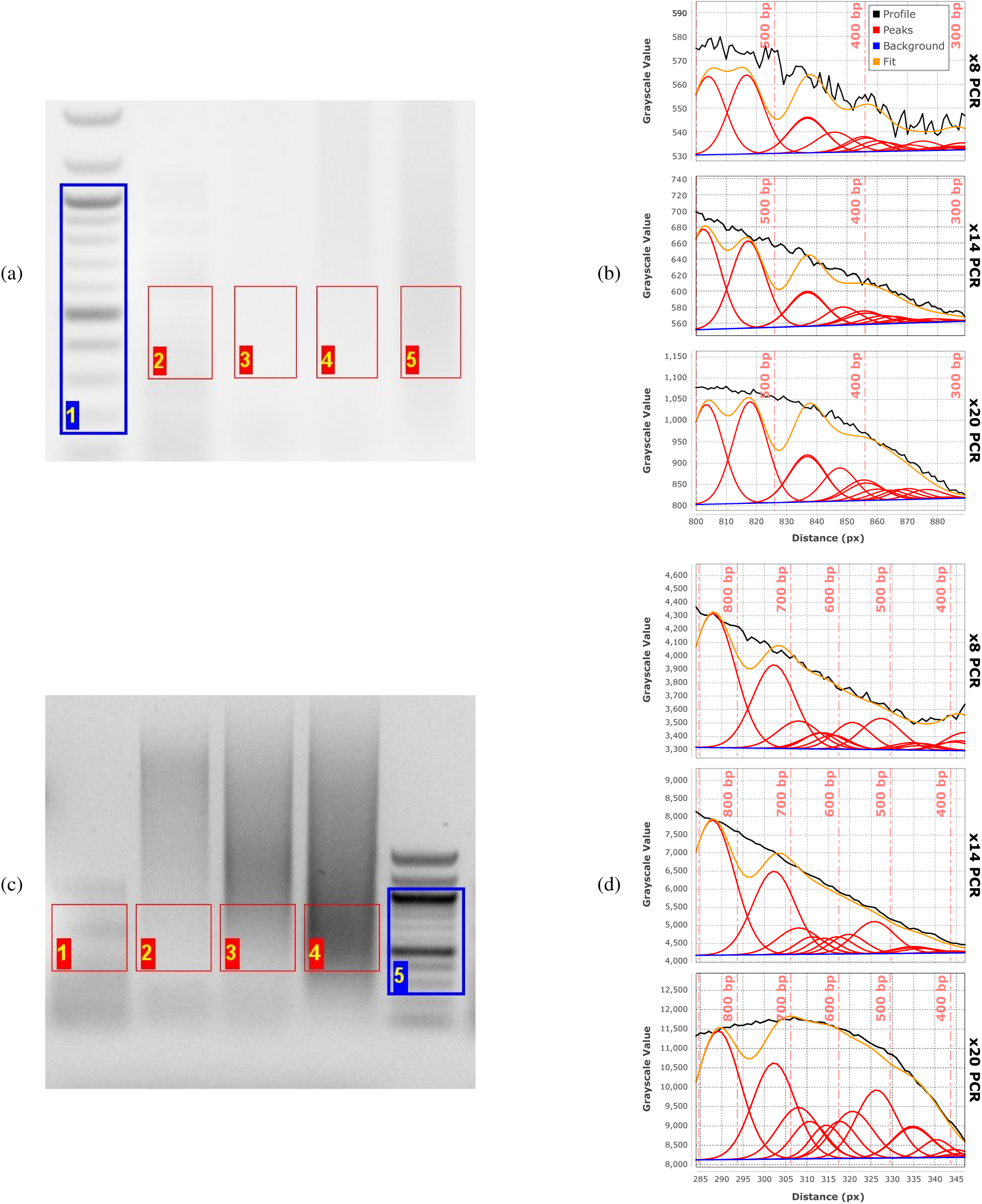
Agarose-gel analysis of tagmentation products of phage-*λ* DNA analyzed on (a) high-resolution and (c) low-resolution (*i.e*., mini) agarose gels. Size-distribution fits for the high-resolution gel (b) and mini gel (d) of the same tagmented *λ*-DNA sample subjected to increasing numbers of PCR-amplification cycles (lanes 3-5 in (a), lanes 2-4 in (c)): 8 cycles (top plots), 14 cycles (middle plots), and 20 cycles (bottom plots) in both (b) and (d). Vertical dashed lines (light red in (b), (d)) give the positions of maxima in the discrete molecular-weight ladder (blue ROI in (b), (b)).

The fragments in an unknown distribution are approximated by a known distribution of fragments having similar electrophoretic mobilities. Figure 2 shows fits to continuous profiles for an unknown fragment-size distribution (a tagmentation reaction on bacteriophage-*λ* DNA, Lanes 3-5 in 2a and Lanes 2-4 in 2c) using the *λ*-*Aci*I digest, a known fragment-size distribution, as a model (Lane 2 in 2a and Lane 1 in 2c). As expected, the algorithm performs best in the range of fragment sizes that are well covered by the known distribution, but produces lower-quality fits in size ranges where there is limited coverage by the *λ*-*Aci*I digest, (*i.e., >* 425 bp).

### Sensitivity to gel resolution

We carried out a pilot assessment of utility of the plug-in by analyzing apparent fragment-size distributions for samples subject to distinct PCR amplification protocols and electrophoretic-separation conditions. In particular, we were interested in assessing whether profiling of gel-electrophoresis patterns using our tool could anticipate differences in size distributions observed in high-throughput Illumina DNA-sequencing experiments for different samples and conditions. Working with a set of samples amplified through different numbers of PCR cycles, we compared plug-in and sequencing output from (*i*) long-format, high-resolution agarose gels (Figure 2a, 2b), and (*ii*) short-format, lower-resolution (minigel) electrophoresis systems (Figure 2c, 2d). Identical samples were run at the same field strength (3.0 V cm-1) on both gels, but for different durations: 16 h in the case of the long-format gel and 2 h in the case of the minigel. Standards consisted of the Hi-Lo ladder, 100-bp ladder, and and the *λ*-*Aci*I digest; experimental lanes contained products of a tagmentation reaction with phage *λ* genomic DNA that were subjected to differing numbers of PCR-amplification cycles.

The effect of sub-optimal fragment separation on minigels is apparent when we compare electrophoretic fragment-size distributions obtained after 8, 14, and 20 PCR cycles (Figure 3). In particular, between 14 and 20 cycles there is a substantial increase in the subpopulation of fragments less than 500 bp in size. The effect of size-distribution distortion is especially prominent in the minigel results and, because of the relatively small region being excised, more difficult to mitigate through selective excision of an appropriate region of the gel. These results both confirm the utility of the plug-in analysis package and highlight the extent to which gel resolution has a significant effect on NGS-library characterization.

**Figure 3.**
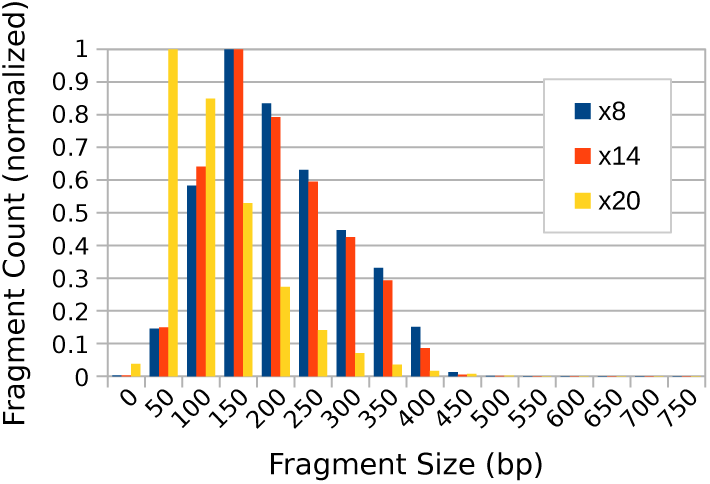
Fragment-library size distributions obtained from bioinformatic analysis of MiSeq sequencing output. A tagmented sample of phage *λ* genomic DNA was subjected to 8, 14 and 20 cycles of PCR amplification, respectively. The fragment counts in each bin are normalized with respect to the maximum value in each distribution.

Further evaluation using the plug-in examined differences in size distributions obtained both from analysis of the lane profiles and from MiSeq post-sequencing fragment-size distributions for the completed library (see Sequencing protocol and bioinformatic analysis). Although input- and output-size profiles for the MiSeq will show some distortion due to the known preference for smaller fragments during bridge amplification, measurement of input material with the plug-in allows prediction of conditions most likely to alter that pattern of eventual fragment densities. Figure 3 quantitatively shows the effect of increasing numbers of amplification cycles on the fragment-size distributions obtained from MiSeq output. As with the data from the plug-in analysis, there is a clear increase in low-molecular-weight bias between 14 and 20 amplification cycles. Although additional aspects of library-preparation and sequencing workflow may contribute to biases in the apparent molecular-weight distributions, it is clear that libraries prepared using limited PCR amplification and careful size selection on high-resolution agarose gels are likely to provide higher fidelity sequencing output.

### Sensitivity to image-acquisition hardware

The images for the gel in Figure 2a were acquired using two different cameras (C1 and C2 in Table 1), whereas images for the gel in Figure 2c were acquired using all four camera systems. Figures 4 and S1 compare the values of average fragment size estimated using the plug-in for both the mini- and high-resolution gel data (numerical data for Figure 4 can be found in Tables S2 and S1). Each data set reports the apparent average fragment size for the *λ*-*Aci*I digest and for different numbers of PCR-amplification cycles of a tagmentation reaction analyzed on a single gel, but imaged using different hardware. The error bars show the standard deviation above and below the mean, *±*1*σ*_*d*_, of the fitted distribution of fragments, as a measure of the distribution’s spread. In general, the differences are modest with the largest discrepancies occurring between high-resolution scientific cameras having appropriate bandpass optical filters and simpler camera systems (such as cell-phone cameras), which generate uncalibrated RGB images. We speculate that the absence of a bandpass filter alters the spectral content of the captured signal, which in the absence of calibration, may report inaccurate signal intensities upon conversion from RGB to grayscale.

**Figure 4.**
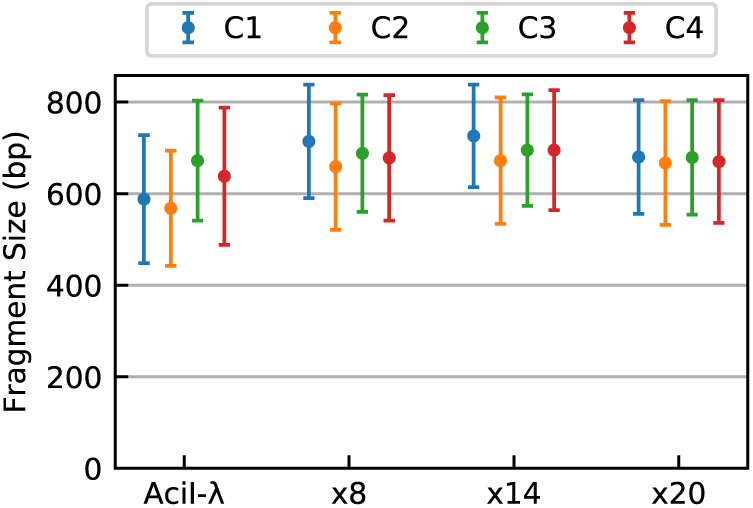
Dependence of plug-in output on camera hardware. Estimates of average fragment size obtained for images of the low-resolution gel (Figure 2c collected using four different camera systems, C1-C4 (Table 1). The plug-in measurements of average fragment size were compared for the known *λ*-*Aci*I digest and the same *λ*-genome tagmented samples analyzed in Figure 2. Error bars indicate the standard deviation (*i.e*.,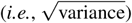) of the fitted distribution. Numerical data for this experiment are also provided in Table S2.

To further test the robustness of our approach, we processed a gel image of tagmentation products taken at different exposure levels. The ROIs in Figure 6a, 6b show the excised regions from which libraries were prepared for this analysis. As shown in Figure 6c, 6d, the apparent average fragment sizes are independent of the imaging exposure time. Detailed results are also reported in Table S3.

### Effect of fragment-size densities in reference distributions

The plug-in provides better profile fits when the reference distribution predominantly consists of DNA fragments in the size range of interest. This is because we generally wish to restrict band widths (FWHM) for both discrete and continuous profiles to values similar to those of co-localized peaks present in the discrete reference ladder. To provide improved profile fits in regions where the reference distribution has fewer peaks, the constraint that the FWHM of an unknown fragment’s peak would be similar to that of a comparably sized fragment in the reference ladder (see Discrete and known continuous patterns for details) has to be relaxed, with the assumption that a single Gaussian peak in the basis set covers a range of fragment sizes that are present in the analyzed distribution.

To examine the effect of limiting reference-peak coverage we created hypothetical reference distributions by regularly removing some basis-set peaks present in the *λ*-*Aci*I digest. Figure 5a shows the gel used for this example and Figure 5b (Table S4) reports the fitted distribution’s mean fragment size and standard deviation. Table S5 reports the absolute RMS as a measure of the closeness of fit for the respective cases.

**Figure 5.**
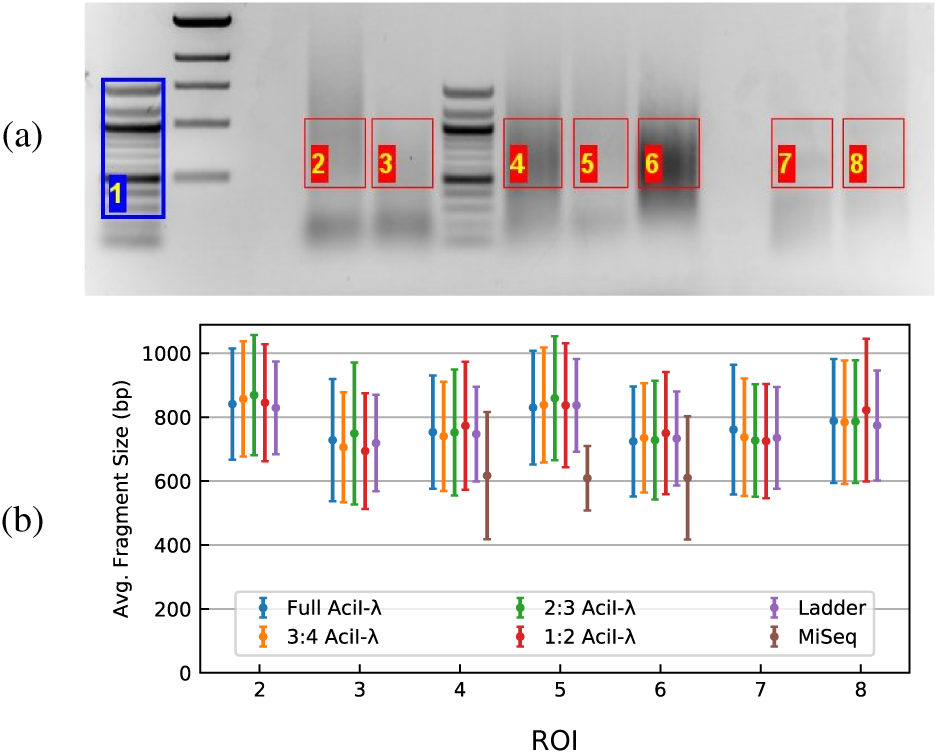
Data used to examine the effect of fragment-density test. (a) The gel was analyzed considering the portions delimited by the ROIs, using lane 1 (blue) as the reference ladder. ROIs 2–6 are tagmentation products of human genomic DNA; 7 and 8 are *λ*-*Aci*I digest samples (b) Summary plot of the fit for the gel in (a). Fragment-size mean and standard-deviation values for model distributions containing varying numbers of reference fragments (peaks). Symbols are colored according to the proportion of the original *λ*-*Aci*I basis fragments retained in the reference distribution; thus, 1:2 means that every other peak was eliminated from the fit, 2:3 every third peak, etc. Data are also reported in Table S4.

### Comparison with TapeStation^®^ output

Tagmented libraries prepared from *C. elegans* genomic DNA were analyzed by agarose-gel electrophoresis as shown in Figure 6 and also using Agilent’s TapeStation^®^ (TS) system. We excised two different sections of the gel from each lane to obtain distinct fragment distributions having different average fragment sizes, as indicated by the respective ROIs. The libraries were reisolated from the gel as described (see Library preparation and gel-electrophoresis-based size selection) and reanalyzed on an Agilent 2200 TapeStation. Here we compare the fragment-size distributions obtained by using the plug-in with those provided by TS output. Generally, fragments of lower molecular size (300-500 bp) are relevant for NGS; however we investigated distributions of larger fragment sizes to determine the effectiveness of our plug-in for other applications. Comparison of plug-in and TS results are shown in Figure 6e and 6f (Table S6).

**Figure 6.**
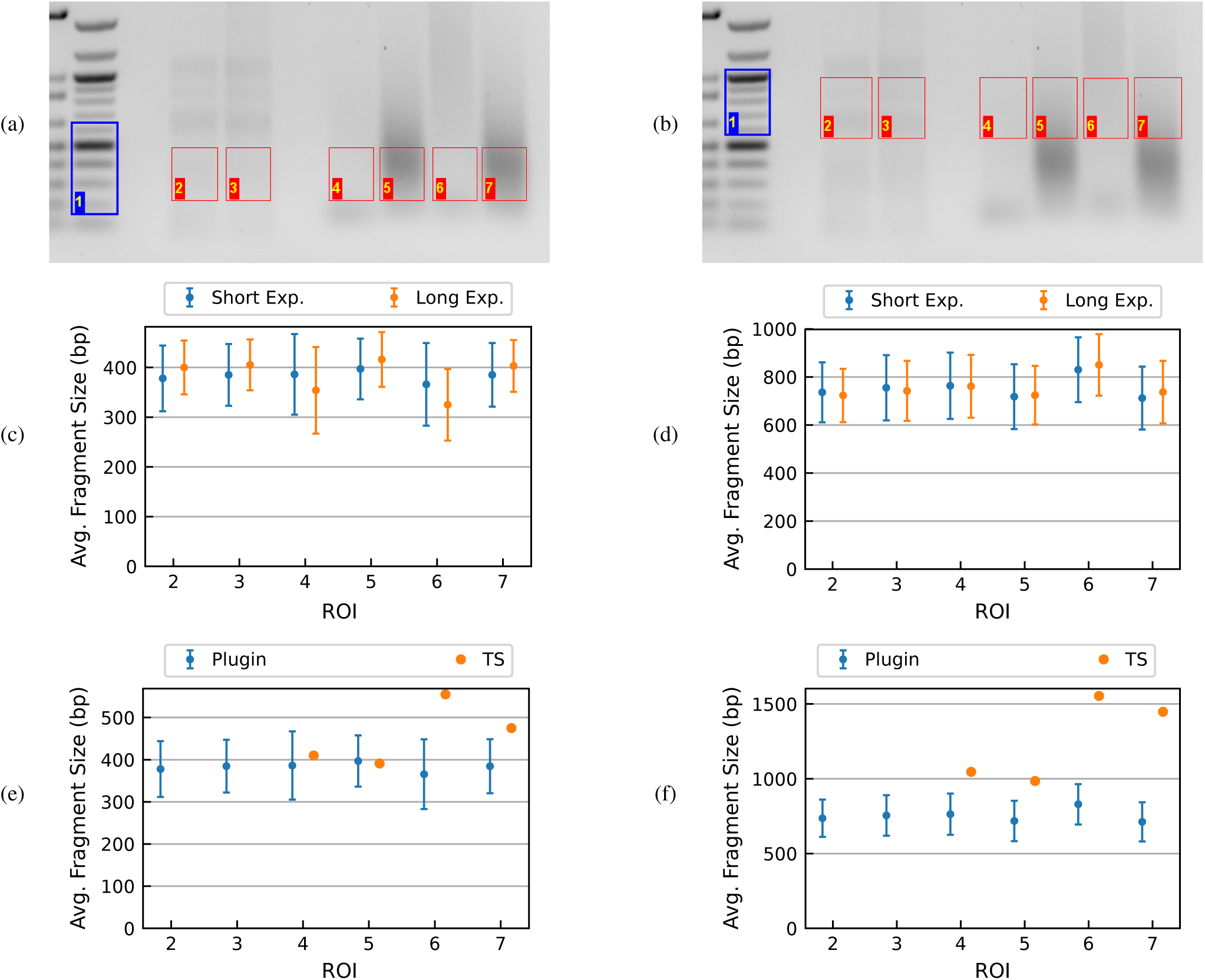
Tagmented libraries prepared from *C. elegans* genomic DNA. Gel images show the sections corresponding to **a** low-molecular-weight and **b** high-molecular-weight fractions. In both cases, lanes 2 and 3 are *λ*-*Aci*I digest samples; lanes 4-7 *C. elegans* genomic-DNA tagmentation reactions amplified by PCR under different conditions. Summary plot of average fragment size based on short- or long-exposure images, **c** low-molecular-weight, and **d** high-molecular-weight fractions (Table S3 in table form). Comparison of average fragment sizes obtained using the plug-in to those generated by TapeStation output for the same DNA fractions: **e** low molecular-weight, and **f** high molecular-weight (Table S6 in table form).

## DISCUSSION

We have developed an ImageJ plug-in for the quantitation of gel-electrophoresis images that can analyze both discrete and continuous gel patterns. This work was motivated by the lack of rigorous and inexpensive DNA size-analysis tools for NGS library preparation. Knowing the distribution of fragment sizes and, hence, the average concentration of DNA termini in a sequencing library is an important quality-control (QC) step, particularly in amplicon-based sequencing.

In the case of discrete patterns, the plug-in can directly quantify relative amounts of DNA based on the intensity and width of the gel bands. In the case of continuous patterns, the plug-in estimates the fragment-size distribution based on a model of superimposed Gaussian peaks derived from a suitable known standard. For NGS library preparation the user specifies, through a rectangular ROI, the region of the gel lane and hence the subset of fragmented products that make up the library. The plug-in provides an estimate of the actual DNA-mass distribution, detailed information that typically goes beyond that provided by specialized QC tools.

We carried out a case study for the determination of library-size distributions using the Nextera^®^ tagmentation protocol. The distribution of fragment sizes in the input library is important in the performance of amplicon-based NGS. Fragments that are too small will be favored during the bridge amplification step, hindering the sequencing of longer fragments; fragments that are too large pose problems with dye localization. Using the plug-in, we were able to quantify a fragment-size bias caused by excessive PCR amplification. Awareness of this amplification bias can improve QC in the preparation of NGS libraries.

The plug-in performs well with both high-resolution and short-format (*i.e*., mini-) gels (see section Image acquisition); however, analysis is most efficient when bands in the reference ladder of DNA standards are well separated and peaks in the reference lane can be detected automatically. Apparent DNA-size distributions were broadly insensitive to image characteristics and quality in our study, which used four different camera systems ranging from scientific CCDs to cell-phone cameras. This shows that the approach implemented by the plug-in can robustly analyze raw image data from a wide variety of sources. From our experience, the precision and accuracy afforded by the approach is more than sufficient for guiding sample preparation in a wide range of sequencing modalities and applications. On-going tool development and applications will investigate effects of digital image compression, conversion from RGB to grayscale, or changes in pixel-value precision when using sensors with different dynamic ranges. For NGS applications, the magnitude of the variations observed in predicted fragment-size averages obtained using different gel cameras are not expected to be propagated in such a way to affect cluster density predictions.

As a final validation step we compared the plug-in’s estimates of average fragment size with that generated by Agilent’s TapeStation^®^ (TS) instrument. The TS system has a list price of approximately $50 000 and significant consumables costs. Moreover, the TS software reports the most-probable fragment size (*i.e*., the mode) along with graphical output of the apparent size distribution. In most cases, the plug-in output is in good agreement with TS output. However, for some of the samples TS reports unreasonable mode values for the size distribution. This happens with very flat profiles where a peak is not detectable. Our approach, while not as ‘hands-off’ as the TS system, can handle lower signal-to-noise line profiles where at least some slope is detectable in the fragment-size distribution.

## ACKNOWLEDGEMENTS

We thank early users whose feedback helped refine the plug-in.

## REFERENCES

1. Steven Head, Kiyomi Komori, Sarah LaMere, Thomas Whisenant, Filip Van Nieuwerburgh, Daniel R Salomon, and Phillip Ordoukhanian. Library construction for next-generation sequencing: Overviews and Challenges. Biotechniques, 56(2):61–, 2014.

2. Rachel Marine, Shawn W. Polson, Jacques Ravel, Graham Hatfull, Daniel Russell, Matthew Sullivan, Fraz Syed, Michael Dumas, and K. Eric Wommack. Evaluation of a transposase protocol for rapid generation of shotgun high-throughput sequencing libraries from nanogram quantities of DNA. Applied and Environmental Microbiology, 77(22):8071–8079, 2011.

3. Nancy C. Stellwagen. Electrophoresis of DNA in agarose gels, polyacrylamide gels and in free solution. Electrophoresis, 30(SUPPL. 1):188–195, 2009.

4. Yasutoshi Shimooka, Jun Ichi Nishikawa, and Takashi Ohyama. Most methylation-susceptible DNA sequences in human embryonic stem cells undergo a change in conformation or flexibility upon methylation. Biochemistry, 52(8):1344–1353, 2013.

5. Stephen D Levene. Analysis of DNA Topoisomers, Knots, and Catenanes by Agarose Gel Electrophoresis. In Duncan J Clarke, editor, DNA Topoisomerases: Methods and Protocols, pages 11–25. Humana Press, Totowa, NJ, 2009.

6. Alexandre A. Vetcher, Abbye E. McEwen, Ramzey Abujarour, Andreas Hanke, and Stephen D. Levene. Gel mobilities of linking-number topoisomers and their dependence on DNA helical repeat and elasticity. Biophysical Chemistry, 148(1-3):104–111, 5 2010.

7. E.M. Southern. Measurement of DNA length by gel electrophoresis. Analytical Biochemistry, 100(2):319–323, 12 1979.

8. J.K. Elder, A. Amos, E.M. Southern, and G.A. Shippey. Measurement of DNA length by gel electrophoresis: I. Improved accuracy of mobility measurements using a digital microdensitometer and computer processing. Analytical Biochemistry, 128(1):223–226, 1 1983.

9. J.K. Elder and E.M. Southern. Measurement of DNA length by gel electrophoresis II: Comparison of methods for relating mobility to fragment length. Analytical Biochemistry, 128(1):227–231, 1 1983.

10. Morgane Boone, Andries DeKoker, and Nico Callewaert. Capturing the ome: the expanding molecular toolbox for RNA and DNA library construction. Nucleic Acids Research, 46(6):2701–2721, 2018.

11. Daniel Aird, Michael G. Ross, Wei Sheng Chen, Maxwell Danielsson, Timothy Fennell, Carsten Russ, David B. Jaffe, Chad Nusbaum, and Andreas Gnirke. Analyzing and minimizing PCR amplification bias in Illumina sequencing libraries. Genome Biology, 12(2):R18, 2011.

12. Margaret A. Taub, Hector Corrada Bravo, and Rafael A. Irizarry. Overcoming bias and systematic errors in next generation sequencing data. Genome Medicine, 2(12), 2010.

13. Massa J Shoura, Idan Gabdank, Loren Hansen, Jason Merker, Jason Gotlib, Stephen D Levene, and Andrew Z Fire. Intricate and Cell Type-Specific Populations of Endogenous Circular DNA (eccDNA) in Caenorhabditis elegans and Homo sapiens. G3: Genes—Genomes—Genetics, 7(10):3295 LP–3303, 10 2017.

14. Michael S. Akhras, Erik Pettersson, Lisa Diamond, Magnus Unemo, Jennifer Okamoto, Ronald W. Davis, and Nader Pourmand. The Sequencing Bead Array (SBA), a Next-Generation Digital Suspension Array. PLoS ONE, 8(10):1–13, 2013.

