## Supplemental Information for "Deconvolution of Nucleic-acid Length Distributions: A Gel Electrophoresis Analysis Tool and Applications"

### SUPPLEMENTARY FIGURES AND TABLES

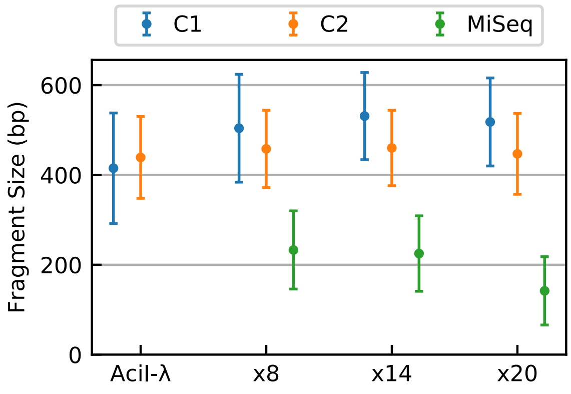


**Figure S1.** Comparison of average DNA-fragment size obtained from plug-in and MiSeq analysis of NGS libraries. NGS data are for phage-$\lambda$ DNA libraries subjected to increasing numbers of PCR-amplification cycles as shown in Figure 2a. The same data are also reported in Table S1.

**Table S1.** Comparison of plug-in output for four camera systems (Table 1) used in acquiring images of the high-resolution gel, shown in Figure 2a. Uncertainty values in Tables are conservatively reported in terms of the standard deviation of the fragment-size distribution, $\pm 1\sigma_d$, as quantified by the plug-in. For this gel-purified sample, *MiSeq*® bioinformatic data are also available.

| | **C1** | **C2** | **MiSeq®** |
|---|---|---|---|
| $\lambda$-*Aci*I Digest | 415±123 | 439±91 | - |
| 8 PCR Cycles | 504±120 | 458±86 | 233±87 |
| 14 PCR Cycles | 531±97 | 460±84 | 225±84 |
| 20 PCR Cycles | 518±98 | 447±90 | 142±76 |

**Table S2.** Comparison of plug-in output for four camera systems (Table 1.) used in acquiring images of the mini-gel, shown in Figure 2c. Uncertainty values in Tables are conservatively reported in terms of the standard deviation of the fragment-size distribution, $\pm 1\sigma_d$, as quantified by the plug-in.

| | **C1** | **C2** | **C3** | **C4** |
|---|---|---|---|---|
| $\lambda$-*Aci*I Digest | 588±140 | 568±126 | 672±131 | 638±150 |
| 8 PCR Cycles | 714±124 | 659±138 | 688±128 | 678±137 |
| 14 PCR Cycles | 726±112 | 672±138 | 695±122 | 695±131 |
| 20 PCR Cycles | 680±124 | 667±135 | 679±125 | 670±134 |

**Table S3.** Tabular data for the plot in Figure 6b, 6c. Size distribution average $\pm 1\sigma_d$, standard deviation. The estimated average fragment size for the $\lambda$-*Aci*I sample is 329 and 734 bp for the *low*-MW and *high*-MW fractions, respectively.

| **ROI** | **Low Cut** | | **High Cut** | |
|---|---|---|---|---|
| | **Low Ex.** | **High Ex.** | **Low Ex.** | **High Ex.** |
| 2 | 378±66 | 400±54 | 736±125 | 723±111 |
| 3 | 385±62 | 405±51 | 755±136 | 742±125 |
| 4 | 386±81 | 354±87 | 763±138 | 761±131 |
| 5 | 397±61 | 416±55 | 718±135 | 724±122 |
| 6 | 366±83 | 325±72 | 830±135 | 850±128 |
| 7 | 385±64 | 403±52 | 712±131 | 737±130 |

**Table S4.** Comparison of the distributions' average fragment size calculations when using different subsets of the reference fragment distribution, the $\lambda$-*Aci*I digest. Size distribution average $\pm 1\sigma_d$, standard deviation.

| ROI | Full $\lambda$-*Aci*I | 3:4 $\lambda$-*Aci*I | 2:3 $\lambda$-*Aci*I | 1:2 $\lambda$-*Aci*I | Ladder | MiSeq® |
|---|---|---|---|---|---|---|
| 2 | 841±174 | 857±180 | 869±188 | 845±183 | 829±145 | - |
| 3 | 728±191 | 706±172 | 749±222 | 694±181 | 719±151 | - |
| 4 | 753±177 | 740±171 | 752±197 | 773±200 | 747±148 | 617±199 |
| 5 | 830±178 | 838±180 | 859±194 | 837±194 | 837±145 | 609±101 |
| 6 | 724±172 | 735±171 | 728±186 | 750±191 | 733±147 | 610±193 |
| 7 | 761±203 | 737±184 | 727±176 | 725±179 | 735±159 | - |
| 8 | 788±194 | 784±193 | 786±192 | 822±223 | 774±172 | - |

**Table S5.** Profile fit accuracy as absolute RMS for the fits for the gel in Figure 5a

| ROI | Full $\lambda$-*Aci*I | 3:4 $\lambda$-*Aci*I | 2:3 $\lambda$-*Aci*I | 1:2 $\lambda$-*Aci*I | Ladder |
|---|---|---|---|---|---|
| 1 | 467.6 | 467.6 | 467.6 | 467.6 | 461.5 |
| 2 | 179.7 | 119.7 | 109.9 | 212.1 | 641.2 |
| 3 | 74.1 | 94.3 | 85.6 | 132.5 | 662.9 |
| 4 | 160.4 | 27.2 | 65.6 | 201.6 | 111.9 |
| 5 | 140.5 | 57.2 | 101.2 | 169.4 | 673.6 |
| 6 | 188.7 | 27.9 | 142.8 | 159.7 | 108.4 |
| 7 | 121.1 | 190.9 | 48.2 | 98.4 | 482.8 |
| 8 | 17.8 | 17.9 | 72.9 | 46.0 | 124.0 |

**Table S6.** Plug-in output compared with TapeStation results. Size-distribution average $\pm 1\sigma_d$, standard deviation, for the plugin output. The calculated average for the $\lambda$-*Aci*I sample is 398 and 871 bp for the *low* and *high* gel portions respectively. The labels in the 2nd and 5th columns, as well as the TS sizes, refer to the plots in section TapeStation Output.

| ROI | | Low Cut | | | High Cut | |
|---|---|---|---|---|---|---|
| | | Plugin | TS | | Plugin | TS |
| 2 | $\lambda$-*Aci*I | 378±66 | - | $\lambda$-*Aci*I | 736±125 | - |
| 3 | $\lambda$-*Aci*I | 385±62 | - | $\lambda$-*Aci*I | 755±136 | - |
| 4 | F1:5 | 386±81 | 410 | B1:1 | 763±138 | 1046 |
| 5 | G1:6 | 397±61 | 391 | C1:2 | 718±135 | 985 |
| 6 | H1:7 | 366±83 | 555 | D1:3 | 830±135 | 1554 |
| 7 | A2:8 | 385±64 | 475 | E1:4 | 712±131 | 1448 |

### TAPESTATION OUTPUT

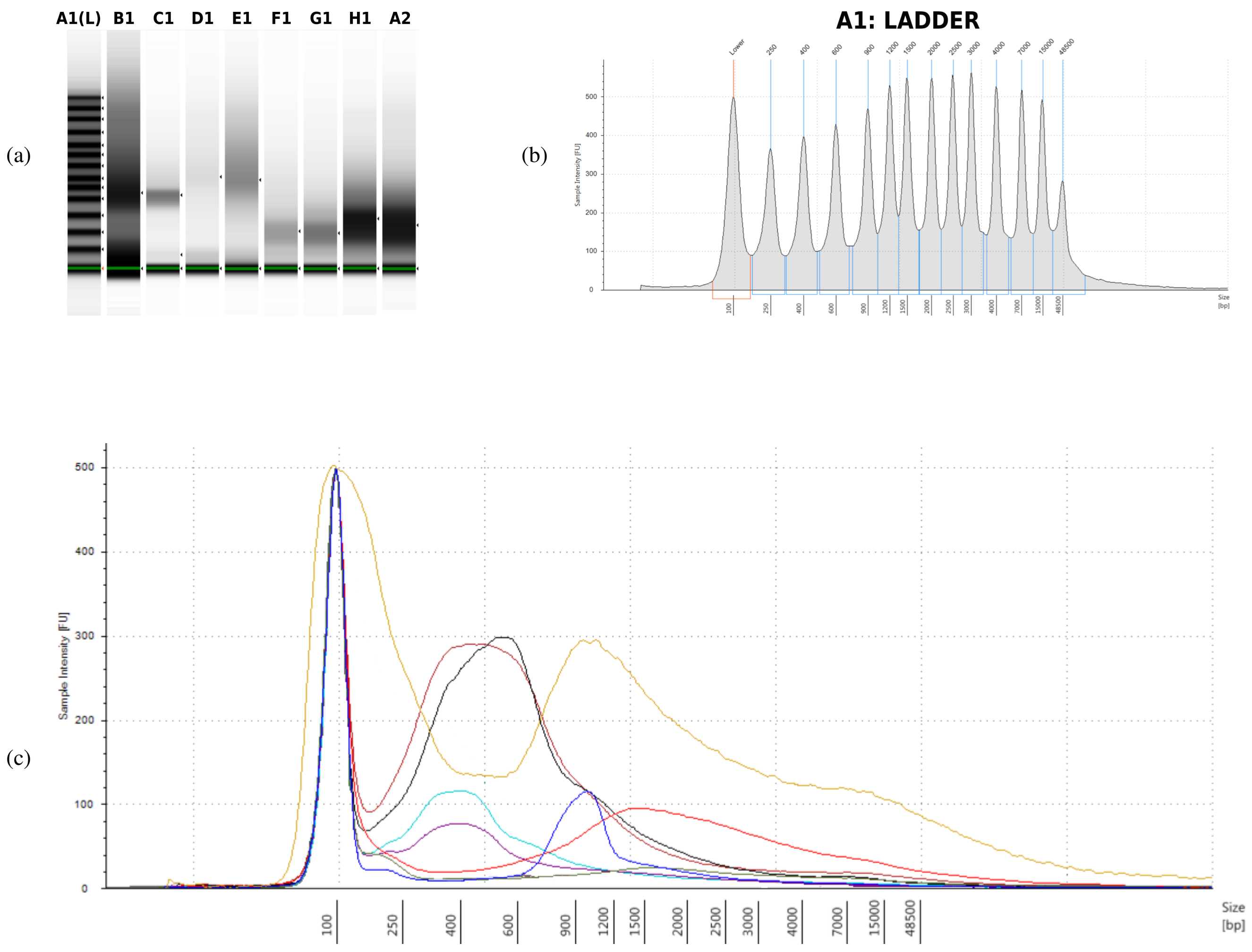


| Well ID | Size (bp) | Well ID | Size (bp) | Well ID | Size (bp) | Well ID | Size (bp) |
|---|---|---|---|---|---|---|---|
| B1 | 1046 | D1 | 1554 | F1 | 410 | H1 | 555 |
| C1 | 985 | E1 | 1448 | G1 | 391 | A2 | 475 |

**Figure S2.** TapeStation® capillary-electrophoretic analysis of tagmented libraries prepared from *C. elegans* genomic DNA. Standard agarose-gel characterization of these libraries is shown in Figure 6. **(a)** original sample profile from the TS, **(b)** the line profile for the ladder of standards ((a), A1(L)) with the location of the peaks; **(c)** line profiles of the analyzed samples and respective main peak location (bp).
